# The bispectral EEG (BSEEG) method quantifies post-operative delirium-like states in young and aged mice after head mount implantation surgery

**DOI:** 10.1101/2024.02.17.580752

**Authors:** Tsuyoshi Nishiguchi, Kazuki Shibata, Kyosuke Yamanishi, Mia Nicole Dittrich, Noah Yuki Islam, Shivani Patel, Nathan James Phuong, Pedro S. Marra, Johnny R. Malicoat, Tomoteru Seki, Yoshitaka Nishizawa, Takehiko Yamanashi, Masaaki Iwata, Gen Shinozaki

## Abstract

Delirium, a syndrome characterized by an acute change in attention, awareness, and cognition, is commonly observed in older adults and has multiple potential triggers, including illness, drug, trauma, and surgery. There are few quantitative monitoring methods in clinical settings. We developed the bispectral electroencephalography (BSEEG) method in clinical research that can detect the presence of and quantify the severity of delirium using a novel algorithm. In the pre-clinical model, we reported that the BSEEG method can capture a delirium-like state in mice following LPS administration. However, its application to post-operative delirium (POD) has not yet been validated in animal experiments. Therefore, this study aimed to create a POD model mouse with the BSEEG method by monitoring BSEEG scores after EEG head-mount implantation surgery throughout the recovery phase. We compared the BSEEG scores of C57BL/6J young (2-3 months old) with aged (18-19 months old) mice for quantitative evaluation of the delirium-like state after the surgery. Postoperatively, both groups showed increased BSEEG scores and a loss of regular diurnal changes in BSEEG scores every daytime and night. In young mice, BSEEG scores and regular diurnal changes recovered relatively quickly to baseline by around postoperative day 3. On the other hand, aged mice had prolonged increases in postoperative BSEEG scores and it reached steady state only after around postoperative day 8. This study suggests the BSEEG method can be utilized to quantitatively evaluate POD and also assess the effect of aging on recovery from POD in pre-clinical model.

## Introduction

Delirium is an acute state of brain failure marked by a sudden onset and a fluctuating course of confusion, inattention, and an abnormal level of consciousness (*1*). Although delirium tends to be transient and self-limiting in some cases, it may persist until hospital discharge or later in others (*2*). Delirium is also a risk factor for poor outcomes, such as high mortality, various in-hospital complications, extended length of hospital stays, institutionalization after discharge, cognitive and functional declines, and dementia (*3, 4*). Moreover, postoperative delirium (POD) is life-threatening in older adults and is the most common postoperative complication in this age group (*5*). Although multiple etiologies have been suggested for delirium (*6*), its pathological mechanism is still not understood. Current diagnosis and treatment of delirium are based primarily on clinical observation and expert opinion because there are few biological-quantitative monitoring methods (*7*). Moreover, delirium is reported to remain undiagnosed in roughly half or more clinical cases (*8*). Thus, despite significant and detrimental consequences to patients’ health and quality of life, there is insufficient research on underlying mechanisms and potential treatments for delirium.

Therefore, an urgent need is to develop an approach to detect delirium and objectively quantify its severity. Our group has been developing a new approach to identify delirium and predict patient outcomes using a novel bispectral electroencephalogram (BSEEG) method that captures the slow wave characteristic of delirium (*9*).

The BSEEG method is based on a simplified and lightweight version of a conventional electroencephalography (EEG) machine that uses only one-channel EEG recording. The BSEEG score can be used to screen for delirium and is also significantly associated with pertinent clinical outcomes, including mortality, length of hospital stay, and disposition after hospital discharge (*10, 11*). The effectiveness of the BSEEG method has been validated in over 1,000 patients (*12*).

Based on the success of the BSEEG methods for human patients in the clinical setting, we pivoted to preclinical studies to lay the groundwork for clarifying delirium pathophysiology to develop effective preventions and treatments (*13*). In preclinical studies of delirium, it is known that a surgical intervention (*14*) or injection of lipopolysaccharide (LPS) (*15*) causes systemic inflammation and neuro-inflammation, leading to a delirium-like state. Following those studies, we showed that the BSEEG method applied to mice could quantify neuroinflammation caused by systemic inflammation induced by intraperitoneal LPS injection (*16*). That report indicates that the BSEEG method applied to an animal model is as useful in detecting abnormal BSEEG scores consistent with delirium as in humans. Thus, as a next step, we tested the validity of the BSEEG method in quantifying surgery-induced neuroinflammation as a model of an animal POD-like state.

Many previous studies have tried verifying POD-like states in rodent models by behavioral tests or tissue imaging to establish a clinically relevant mouse model of POD (*17, 18*). However, these assessments take time and are labor-intensive. Additionally, they are limited to providing insight into a short time window.

Several groups have reported that EEG can help objectively assess responses to LPS and other inflammatory triggers (*19–21*). However, their studies focused on only young mice and did not evaluate the influence of age, which is a well-known major risk factor for delirium. Also, continuous monitoring for quantifying POD-like states of mice has hardly been reported. One paper monitored aged mice in an ICU-like condition with EEG and reported that their EEG and circadian changes showed frequent arousals and slow wave ratios relative to controls, similar to delirious ICU patients (*22*). However, there has not yet been a study monitoring long postoperative periods with EEG and comparing young and aged mice.

To address the issues above, we applied the BSEEG method after an EEG head mount surgery to continuously monitor and quantify BSEEG response from surgical insults. In the present study comparing young and aged mice, we report the differences in BSEEG scores between different age groups after an EEG head mount surgery.

## Methods and materials

### Animals and housing

All male C57BL/6J mice (young: 2–3 months and aged: 18-19months) were purchased from Jackson Laboratory. All mice were housed in the animal housing facility at Stanford University. 4-5 mice were housed in an Innocage^®^ before surgery, with food and water available freely, and each mouse was housed separately after surgery. The facility maintained the housing environments in a 12:12 h light-dark cycle and at a proper temperature and humidity. The animal experiments were conducted following a protocol approved by an Institutional Animal Care and Use Committee (IACUC). At Stanford, the IACUC is known as Stanford’s Administrative Panel on Laboratory Animal Care (APLAC). The APLAC is accredited by the Association for Assessment and Accreditation of Laboratory Animal Care (AAALAC).

### Experimental procedure

1. The schedule of the experiment: EEG recording began immediately after the EEG electrode head mount placement surgery. Postoperative day (PO-Day) ranges from 0 to 14, with day 0 as the surgery day. We also compared the BSEEG time course between young mice (n = 12) and aged mice (n = 15). The recorded data was analyzed through our group’s publicly available web tool (*16*).
2. EEG electrode head mount placement surgery: EEG electrode head mount placement surgery was conducted as in previous studies (*16, 23, 24*). The skull was exposed under isoflurane (1-3% inhalation) anesthesia. The head mount (#8201; Pinnacle Technology, Inc., Lawrence, KS) was attached with its center at the midline of the skull, with its anterior holes at 2 mm anterior to bregma and its posterior holes at 2 mm anterior to lambda ± 2 mm. Four holes were bored into the skull with a 23-gauge needle through holes of the head mount. Four screws (anterior: #8209, posterior: #8212; Pinnacle Technology, Inc., Lawrence, KS) were attached to the skull. Finally, the head mount was fixed with dental cement **(Fig. 1)**. Mice received postoperative analgesia with meloxicam 5mg/kg in subcutaneous injection. After that, mice were connected by wire to an EEG system (#8200-K1-SL; Pinnacle Technology, Inc., Lawrence, KS) and kept connected during a 2-week recovery period **(Fig. 1)**. EEG were recorded from a right frontal cortex using the right anterior screw attached to the skull as EEG2, and a parietal cortex using the right posterior screw attached to the skull as EEG1. A left frontal cortex using the left anterior screw was attached to the skull as a ground and a left parietal cortex using the left posterior screw was attached to the skull as a reference. We analyzed signals from the electrode EEG2 lead for the BSEEG scoring because EEG2 is more sensitive than EEG1 according to the previous study(*16*).

**Figure 1.**
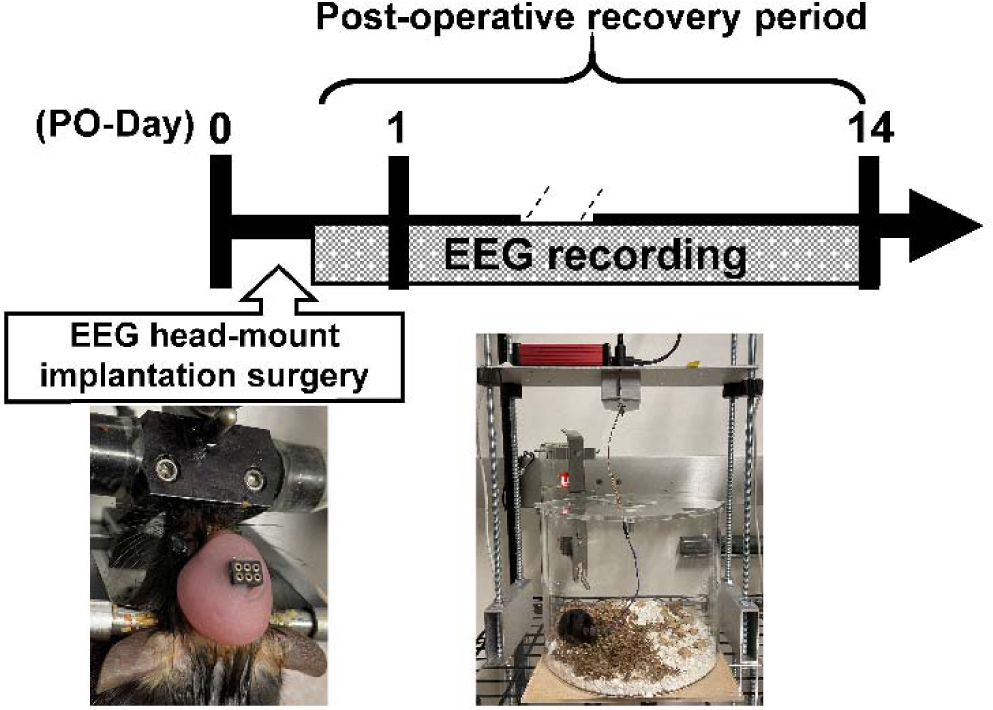
Schematic diagram of the experiment.

### Calculation of BSEEG scores

1. EEG signal processing for BSEEG scores: As raw EEG recordings were digitally converted into power spectral density to calculate BSEEG scores in our previous studies (*9, 25, 26*), animal EEG recordings were analyzed using a similar algorithm to the human study. We used the software SleepPro (Sirenia EEG system; Pinnacle Technology, Inc., Lawrence, KS) to export raw data after Fast Fourier Transformation. We processed it through our algorithm to calculate BSEEG scores. Our BSEEG scores were generated by calculating the ratio between 3 Hz power to 10 Hz power.
2. Web-based BSEEG score calculator: According to a previous study (*16*), we calculated BSEEG scores. After recording raw EEG signals, the data was exported as an EDF file by the default software from the Pinnacle Sirenia EEG system. Then, the EDF file was uploaded to our web-based tool (https://sleephr.jp/gen/mice7days/ ), and every BSEEG score per hour was calculated.
3. BSEEG score: In this study, we defined a BSEEG score as every daytime/night 12 hours average BSEEG score from 7 am/7 pm to highlight regular diurnal changes seen every day over 24 hours as reported by the previous study(*16*). BSEEG scores were plotted on the y-axis, whereas the recording times were displayed on the x-axis. Daytime/night was shown as yellow/gray lines on the background of the x-axis.
4. BSEEG score standardized by the BSEEG score from PO-Day 14 daytime (sBSEEG score): To standardize the degree of differences in the BSEEG scores compared to the BSEEG score when it reaches to its steady state after recovery from surgery, we defined the BSEEG score on PO-Day 14 daytime as sBSEEG = 0. We took this approach because it is not possible to measure the baseline EEG recordings prior to head mount surgery, and because it is known that as general animal postoperative care, animals are fully recovered from 7 to 14 days following surgery(*27*), and thus most mice were estimated to fully recover and reach steady-state of their BSEEG score by PO-Day 14. The BSEEG score standardized by the BSEEG score from PO-Day 14 daytime (sBSEEG score) was defined as the difference between BSEEG scores on each PO-Day daytime/night and BSEEG scores on PO-Day 14 daytime. The sBSEEG scores were plotted as mentioned above. Daytime/night was also shown as mentioned above.

### Statistical analysis

To compare the BSEEG scores or sBSEEG scores between young and aged mice, or those between PO-Day 14 and other PO-Day, a t-test was performed using GraphPad Prism version 9.5.0 for Windows, GraphPad Software (www.graphpad.com). All error bars were shown as means ± standard error of the means (mean ± SEM). P < 0.05 was marked with an asterisk (*) and considered statistically significant.

## Results

The data in Figure 2 shows the results from the 12 young mice **(Fig. 2)**. Each graph represents the postoperative time course of BSEEG scores. Most of the young mice showed that the BSEEG scores comparing on PO-Day 1 with on PO-Day 14, in other words “the increases in sBSEEG score”, were in ranging from 1 to 4 **(Table 1**. No 1, 2, 4, 6, 7, 9, 10, 11) and loss in diurnal changes for several days following the head mount implantation surgery. A few mice show drastic increases in BSEEG scores on PO-Day 1 (> 10) **(Fig. 2 and Table 1**. No 3, 8). However, there were a few mice almost unaffected in BSEEG scores on PO-Day 1**(Fig. 2 and Table 1**. No 5, 12). It is noteworthy that by PO-Day 14 the daytime BSEEG scores from most mice reached the steady state around 4 **(Fig. 2 and Table 1** . No 1, 2, 3, 4, 5, 6, 7, 8, 9, 11, 12).

**Figure 2.**
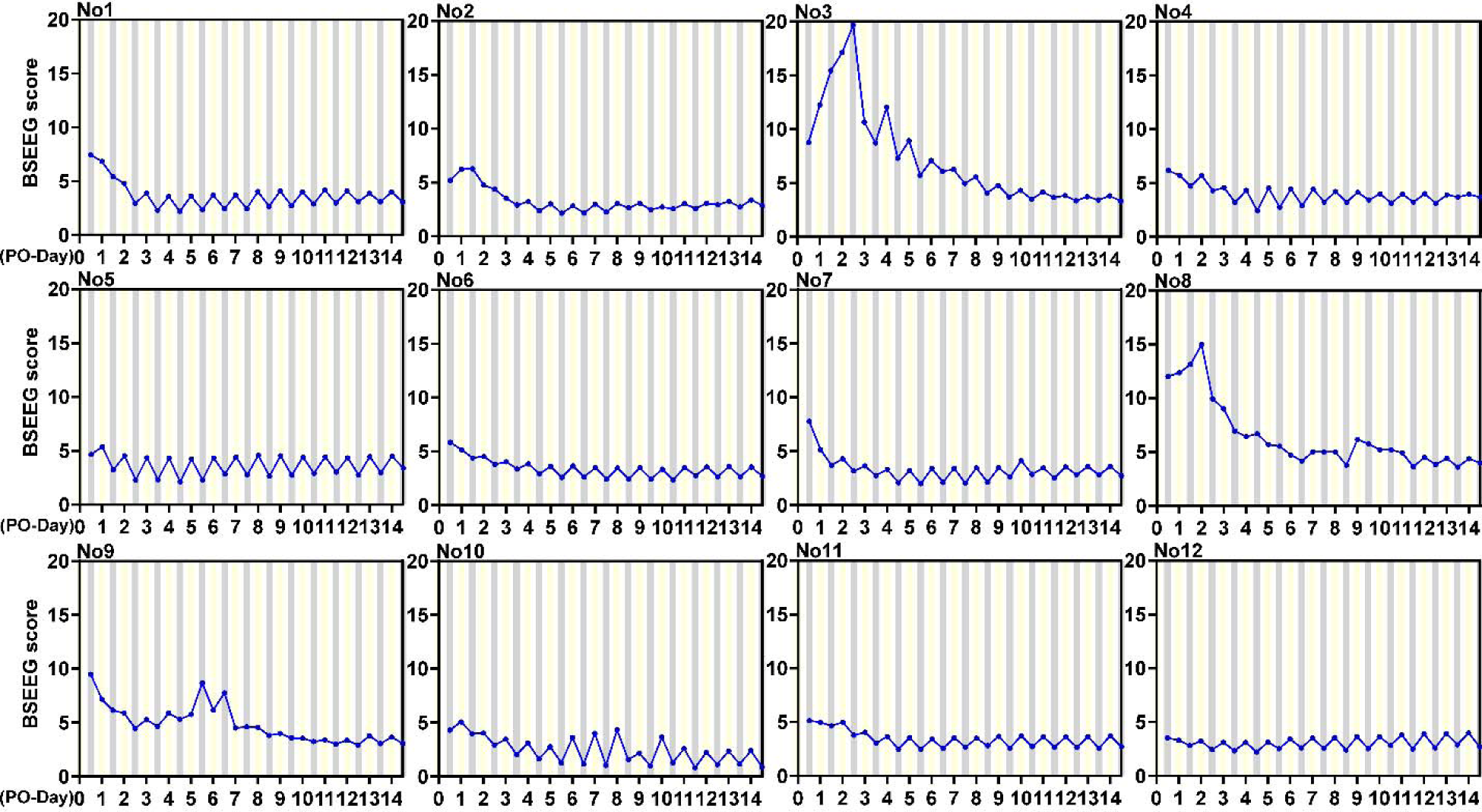
The 14 days of the post-operation time course of each young mouse. BSEEG scores were plotted on the y-axis, whereas the recording times were displayed on the x-axis. Daytime/night was shown as yellow/gray lines on the background of the x-axis.

**Table 1.** Each young mouse’s BSEEG score on PO-Day 1 and 14.

|  | **PO-Day 1**<br>**daytime**<br>**BSEEG score** | **PO-Day 14**<br>**daytime**<br>**BSEEG score** | **PO-Day 1**<br>**daytime**<br>**sBSEEG score** |
| --- | --- | --- | --- |
| **No. 1** | 6.84 | 3.99 | 2.86 |
| **No. 2** | 6.24 | 3.38 | 2.86 |
| **No. 3** | 12.25 | 3.82 | 8.43 |
| **No. 4** | 5.71 | 3.96 | 1.75 |
| **No. 5** | 5.39 | 4.49 | 0.89 |
| **No. 6** | 5.16 | 3.55 | 1.61 |
| **No. 7** | 5.16 | 3.59 | 1.56 |
| **No. 8** | 12.37 | 4.38 | 7.99 |
| **No. 9** | 7.18 | 3.65 | 3.53 |
| **No. 10** | 5.07 | 2.40 | 2.67 |
| **No. 11** | 4.98 | 3.74 | 1.24 |
| **No. 12** | 3.30 | 4.00 | -0.69 |

We combined the data of the post-operative time course from the 12 young mice and calculated the mean BSEEG scores from the night of PO-Day 0 to the night of PO-Day14 **(Fig. 3 and Supplementary Table 1)**. The data points from PO-Day 0 to 3 indicated the recover from the increase in the mean BSEEG scores, which was caused by the effects of the EEG head-mount implantation surgery. The timing of the highest BSEEG was observed at either PO-Day 0 night or PO-Day 1 daytime for most mice **(Fig. 2)**. Regular diurnal changes disappeared during the period between PO-Day 0 and 3 daytime until it returned from PO-Day 3 night **(Fig. 3)**.

**Figure 3.**
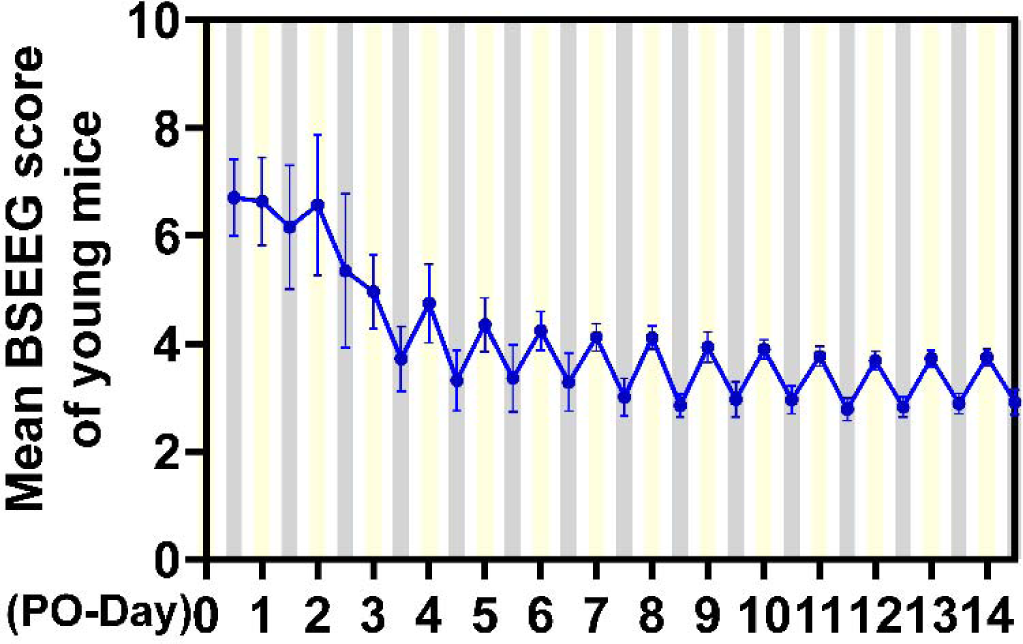
The post-operative time course of the mean BSEEG score of all young mice. BSEEG scores were plotted on the y-axis, whereas the recording times were displayed on the x-axis. Daytime/night was shown as yellow/gray lines on the background of the x-axis. All error bars were shown as means ± standard error of the means (mean ± SEM). P < 0.05 was marked with an asterisk (*) and considered statistically significant.

Fig. 4 shows the results from the 15 aged mice **(Fig. 4)**. These data indicate that the variation in BSEEG scores was more diverse in aged mice than in young mice. Only a few aged mice **(Fig. 4** No 3, 9) showed regular diurnal variation similar to the pattern commonly seen in young mice **(Fig. 2 and Fig. 3)**. However, most other aged mice showed no consistent pattern in the BSEEG scores even at their steady state. They were either flat **(Fig. 4** No 1, 5, 6, 12, 13, 14), reversed diurnal change where BSEEG scores are higher at night **(Fig. 4** No 10, 15), or irregular **(Fig. 4** No 2, 4, 7, 8, 11). A few aged mice showed drastic increases in BSEEG scores (> 10) soon after surgery on PO-Day 1, 2, or 3 **(Fig. 4 and Table 2**. No 4, 6, 10 and 11). The BSEEG scores from many mice showed some recovery from the increase in BSEEG score and the loss of regular diurnal changes from the effect of surgery by PO-Day 14. However, in contrast to young mice, it is remarkable that the BSEEG score patterns observed by PO-Day 14 remained diverse depending on each mouse **(Table 2.)**.

**Figure 4.**
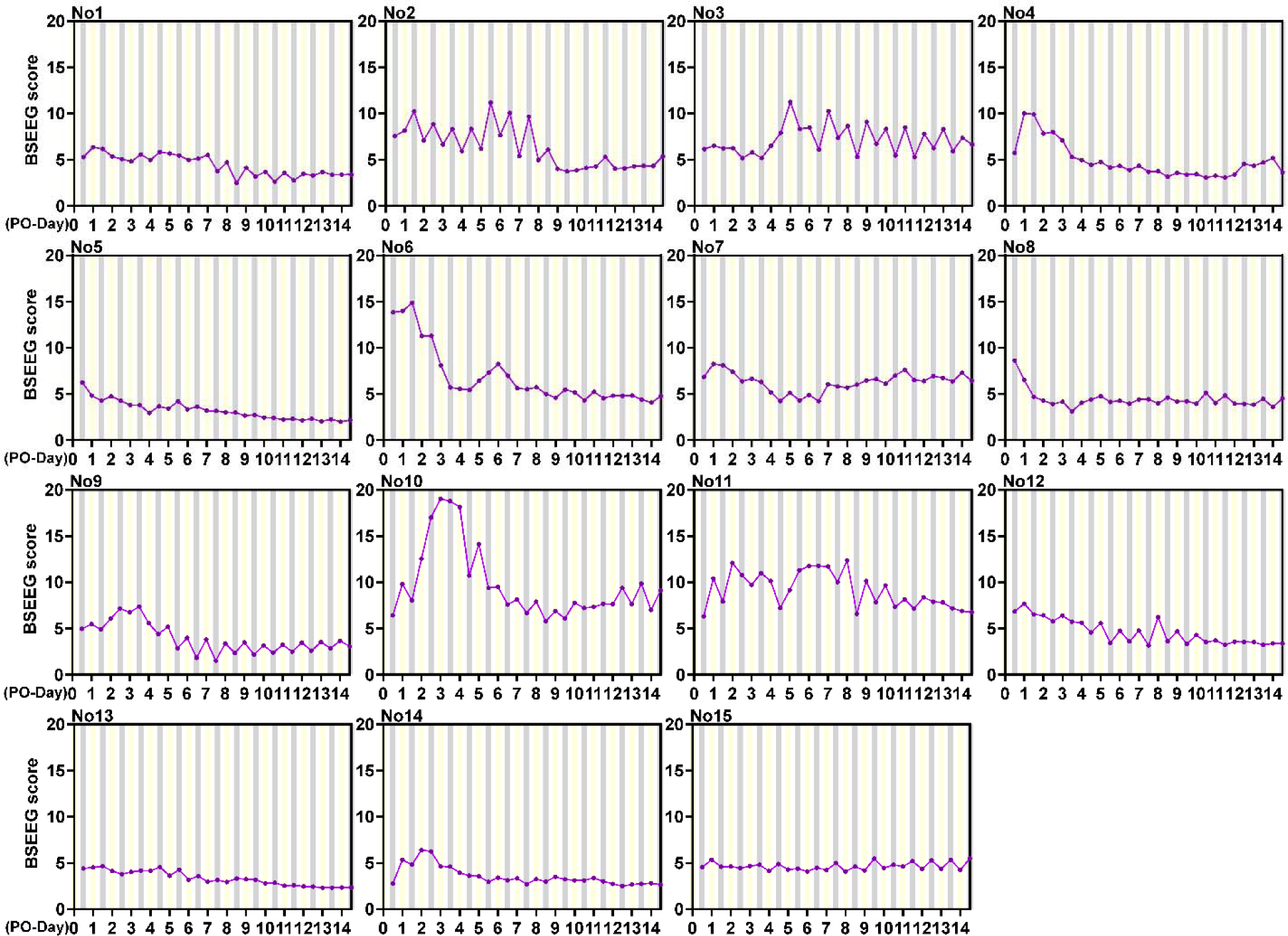
The 14 days of the post-operative time course of each aged mouse. BSEEG scores were plotted on the y-axis, whereas the recording times were displayed on the x-axis. Daytime/night was shown as yellow/gray lines on the background of the x-axis.

**Table 2.** Each aged mouse’s BSEEG score on PO-Day 1 and 14.

|  | **PO-Day 1**<br>**daytime**<br>**BSEEG score** | **PO-Day 14**<br>**daytime**<br>**BSEEG score** | **PO-Day 1**<br>**daytime**<br>**sBSEEG score** |
| --- | --- | --- | --- |
| **No. 1** | 6.35 | 3.39 | 2.96 |
| **No. 2** | 8.15 | 4.31 | 3.84 |
| **No. 3** | 6.51 | 7.36 | -0.85 |
| **No. 4** | 10.02 | 5.19 | 4.83 |
| **No. 5** | 4.81 | 1.99 | 2.82 |
| **No. 6** | 14.00 | 4.07 | 9.93 |
| **No. 7** | 8.25 | 7.30 | 0.95 |
| **No. 8** | 6.50 | 3.59 | 2.91 |
| **No. 9** | 5.47 | 3.65 | 1.82 |
| **No. 10** | 9.80 | 7.01 | 2.79 |
| **No. 11** | 10.41 | 6.90 | 3.51 |
| **No. 12** | 7.69 | 3.38 | 4.30 |
| **No. 13** | 4.52 | 2.35 | 2.17 |
| **No. 14** | 5.33 | 2.81 | 2.53 |
| **No. 15** | 5.34 | 4.25 | 1.09 |

Fig. 5 showed the average BSEEG scores combined from the data of the post-operative time course of the 15 aged mice. The mean BSEEG scores from the night of PO-Day 0 to the night of PO-Day14 were calculated **(Fig. 5 and Supplementary Table 2)**. The data points from PO-Day 0 to around Day 7 remained elevated, indicating the slow recovery from the effects of the EEG head-mount implantation surgery among aged mice **(Fig. 5)**. Overall, there were much less clear diurnal change patterns among aged mice **(Fig. 5)**.

**Figure 5.**
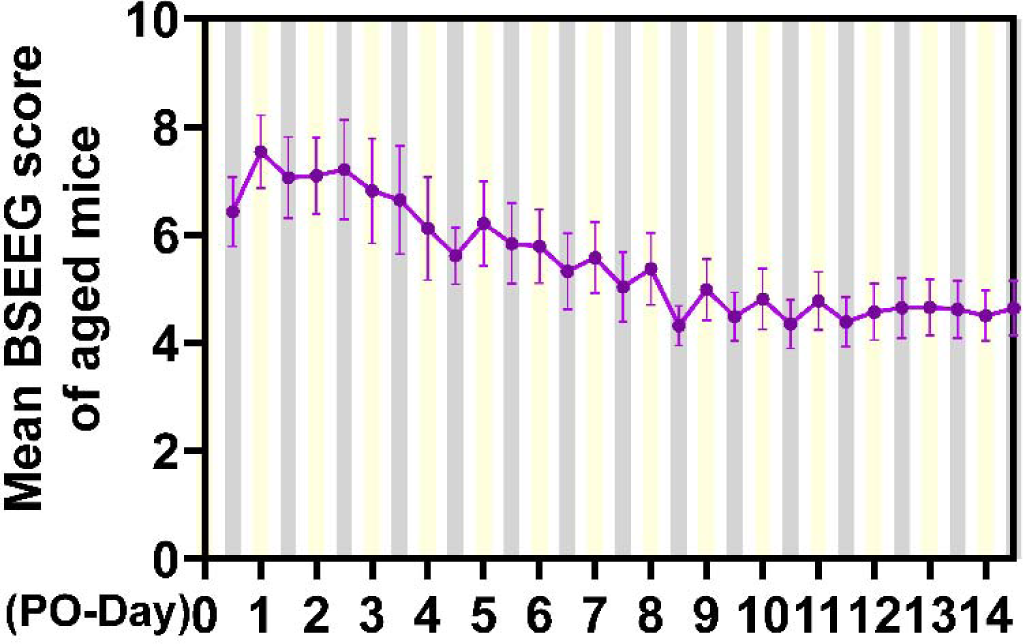
The post-operative time course of the mean of all aged mice. BSEEG scores were plotted on the y-axis, whereas the recording times were displayed on the x-axis. Daytime/night was shown as yellow/gray lines on the background of the x-axis. All error bars were shown as means ± standard error of the means (mean ± SEM). P < 0.05 was marked with an asterisk (*) and considered statistically significant.

Next, we compared the mean BSEEG scores of young and aged mice **(Fig. 6 upper graph and Supplementary Fig. 1)**. Aged mice had significantly higher average BSEEG scores at night than young mice during the whole postoperative period, except from PO-Day 0 to 2 **(Fig. 6 upper graph black asterisks and Supplementary Fig. 1)**. As mentioned previously, the mean BSEEG scores of young mice regained regular diurnal changes early by PO-Day 3 and reached the steady state ranging between approximately 2.5 and 4 **(Fig. 3 and Supplementary Table 1)**. In contrast, the mean BSEEG scores of aged mice did not regain clear diurnal changes and seemed to reach steady state with slight fluctuation around 4.5 to 5 later by PO-Day 9 **(Fig. 5 and Supplementary Table 2)**.

**Figure 6.**
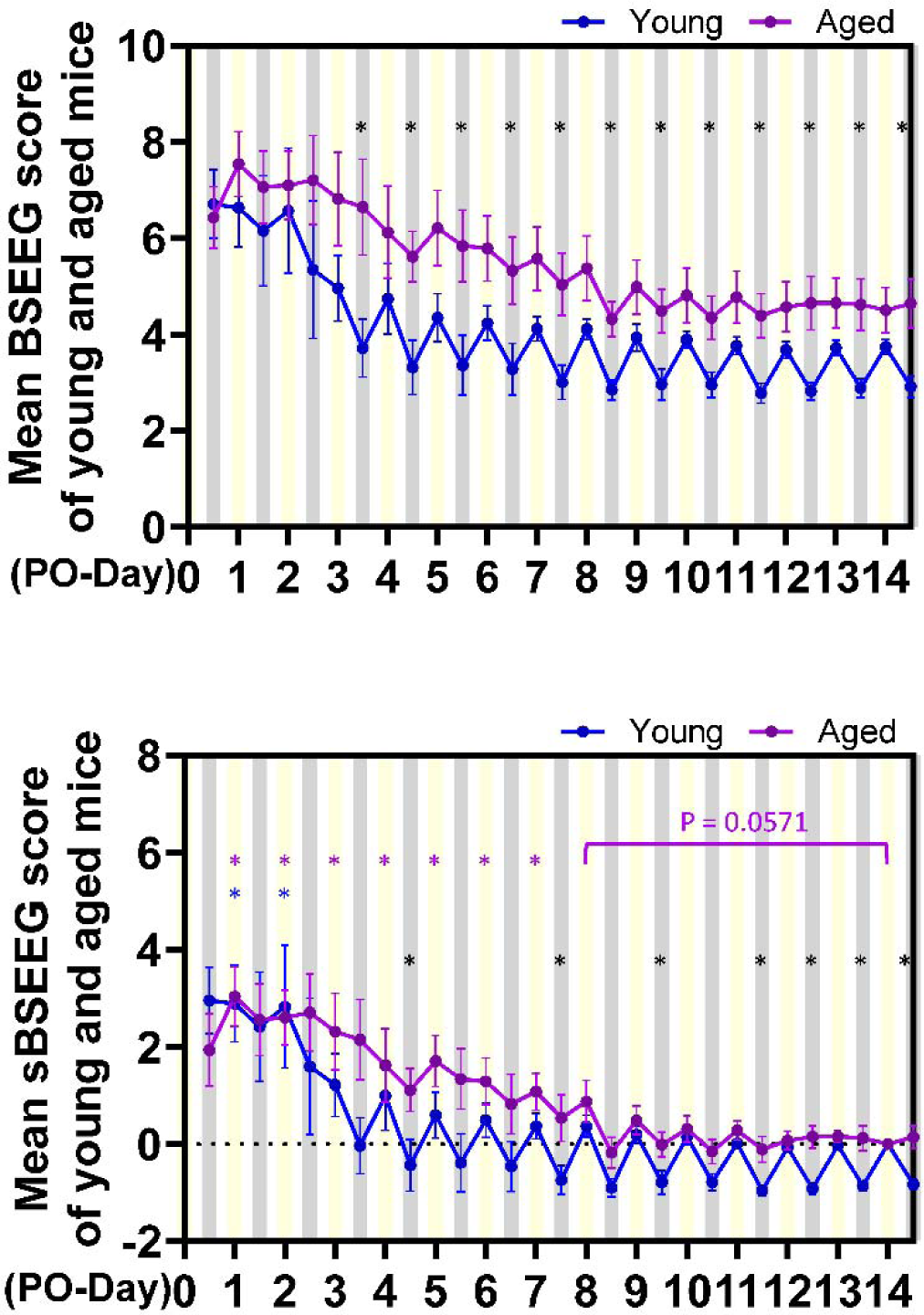
The postoperative time course of the mean BSEEG scores (upper graph) and the mean sBSEEG score (lower graph) in both groups. BSEEG (upper graph) and sBSEEG (lower graph) scores were plotted on the y-axis, whereas the recording times were displayed on the x-axis. Daytime/night was shown as yellow/gray lines on the background of the x-axis. All error bars were shown as means ± standard error of the means (mean ± SEM). P < 0.05 was marked with an asterisk (*) and considered statistically significant. Black asterisks are the significant differences in BSEEG (upper graph) or sBSEEG (lower graph) scores between the young and aged mice. Blue asterisks are the significant difference in sBSEEG scores between the daytime of each PO-Day and PO-Day 14 in the young mice. Purple asterisks are the significant difference in sBSEEG scores between the daytime of each PO-Day and PO-Day 14 daytime in the aged mice.

Because the steady state levels of the mean BSEEG scores were different between young and aged mice, we compared the standardized BSEEG (sBSEEG) scores between young and aged mice **(Fig. 6 lower graph and Supplementary Fig. 2)**. In the young group, there were statistically significant differences in the sBSEEG scores on the daytime of PO-Day 1 and 2 compared with that of PO-Day 14, and there was no statistical significance after the daytime of PO-Day 3 compared with that of PO-Day 14 **(Fig. 6 lower graph blue asterisks and Supplementary Fig. 3)**. On the other hand, in aged mice, there were statistical significances from the daytime of PO-Day 1 to 7 compared with that of PO-Day 14, and the daytime mean sBSEEG scores showed no statistical significance after the daytime of PO-Day 8 compared with that of PO-Day 14 **(Fig. 6 lower graph purple asterisks and Supplementary Fig. 4)**. This graph shows that the value of the postoperative increase on PO-Day 0 to 2 was not significantly different between young and aged mice. The major difference between young and aged mice was the more prolonged duration of the sBSEEG score elevation on PO-Day 3 to 7 in aged mice **(Fig. 6 lower graph)**.

## Discussion

We showed that the BSEEG method, proven effective in detecting delirium in the clinical setting, could also capture and quantify the brain wave response after surgery among young and aged mice. This data supports the usefulness of the BSEEG score in assessing the delirium-like state in postoperative mice. There was a significant difference between young and aged mice in mean BSEEG score following surgery **(Fig. 6, Supplementary Fig. 1)**, as expected based on our previous data showing a more significant response to the same dose of LPS among aged mice than young mice (*16*). In young mice, the effect of surgery on BSEEG score was relatively short: many mice started showing regular diurnal variation soon after surgery, and the BSEEG score recovered to the baseline level by PO-Day 3 **(Fig. 2, 3, and 6 Supplementary Table 1 and 3)**. In aged mice, however, such recovery took longer: diurnal variation in BSEEG scores after surgery was not well established and diverse depending on each mouse, and the BSEEG score did not recover to baseline until PO-Day 8 **(Fig. 4, 5 and 6, Supplementary Table 2 and 4)**.

The prolonged period of elevated BSEEG required to reach a steady state among aged mice is consistent with the fact that older adults are more vulnerable and prone to develop delirium. At the same time, the lack of clear diurnal change even at steady state among aged mice suggested that loss of circadian rhythm may be associated with some unknown mechanisms related to the vulnerability of the older adults to delirium. In fact, recent studies reported the possible role of the circadian timing system in the pathophysiology of delirium(*28, 29*). The circadian timing system is a major determinant of the timing of sleep and sleep structure in humans, and many aspects of sleep vary markedly with circadian phase in both young and older adults(*30*). In general, older adults sleep and wake at earlier times than do young adults, older adults are more likely to report advanced sleep wake phase disorder than are young adults, and older adults are more prone to shift work disorder and jet lag disorder(*31*). One paper reported that circadian rhythm-related factors, decreased REM sleep, lower melatonin levels, and higher cortisol levels, were independently associated with an increased risk of delirium in intensive care unit(*32*). Such vulnerabilities of older adults for delirium may be related to the dysregulation of circadian timing system. Our data showing the irregular diurnal changes of BSEEG score together with its prolonged elevation after surgery among aged mice in this study may support such relationship. Thus, treating dysfunction in circadian timing system may prevent and treat delirium in older adults(*33*). In fact, some studies report that melatonin administration to older adults may represent a potential protective agent against delirium(*34, 35*). The BSEEG method may detect preoperative vulnerability thorough monitoring diurnal changes of BSEEG score and will be expected to evaluate potential preventive or therapeutic effects of the intervention on delirium.

It is well known that the incidence of POD depends on the type of surgery and age (*36*). Under different conditions, the course of the delirium-like state in animal model quantified by the BSEEG method may also differ. While conventional methods for evaluating delirium-like state in mice require behavioral tests at each distinct time point, the BSEEG method allows continuous and quantitative evaluation. Thus, the BSEEG method can have advantages in obtaining more detailed chronological and objective information of the delirium-like state induced under various conditions, including age, sex, surgical type, or drug administration. The BSEEG data from our present study using the postoperative model are consistent with the previous literature. Some studies reported that the impairment of cognitive function and glial activation in young adult animals was observed between PO-Day 1 and 3 but not at PO-Day 7 (*37*) and that the impairment of cognitive function in aged animals was observed later at PO-Day 6 and 7 (*38*). Moreover, one study reported that anesthesia and surgery impair the blood-brain barrier and cognitive function in aged mice more than adult mice (*39*). It is also reported that microglia release more cytokines in the brains of aged animals than in younger ones in response to exogenous insults, indicating that activated microglia may be involved in the pathogenesis of cognitive decline in aged animals (*40, 41*).

To date, various studies have investigated the relationship between delirium and aging, and numerous mechanisms have been reported to underlie the changes in the aging brain and their delirium-related effects (*42*). Pro-inflammatory changes of microglia and macrophages in the immune system (*43*), inadequate blood flow and increased blood-brain barrier permeability in parenchymal vessels (*44*), decreased connectivity in nerve cell systems (*45*), and reduced mitochondrial efficiency (*46*) are among such mechanisms. In addition, our group has been focusing on the influence of age and investigating the potential role of epigenetic mechanisms, especially DNA methylation (DNAm), in the pathophysiological mechanism of delirium (*47, 48*). Our data clearly showed that DNAm in the TNF-alpha gene decreases along with age, together with the age associated increase of the gene expression. The data suggests that aging can make people more prone to be “inflamed” (*40*).

These wide ranges of mechanisms may also be involved as potential factors that cause the differences in BSEEG scores with age in the present study. In other words, identification of modifiable factors to prevent BSEEG increase, especially among the aged group, could lead to novel therapeutic interventions to prevent and treat delirium in humans. To that end, this novel BSEEG method applied to mice can be an important tool for future delirium/POD research.

We acknowledge that this study has several limitations. First, although the BSEEG method was used, we did not investigate its relationship with the behavioral experiments or the pathological examination of brain tissues. Second, we did not investigate the details of individual differences in vulnerability. As individual differences in resiliency and vulnerability in response to the exogenous insults may be based on the core pathophysiological mechanisms of delirium, we need to study these factors more in detail in the future. Third, since the EEG head mount was placed directly into the brain, the location and depth of screw insertion may have affected the obtained EEG data. However, considering the generalized slow waves observed in delirium, it is unlikely that a slight difference in insertion site would be significant enough to interfere with the observed signals. Finally, since the screw is inserted directly into the brain, it causes some level of direct brain injury. As direct brain damage can cause acute brain dysfunction beyond neuroinflammation alone, including energy deprivation and metabolic disturbances (*49*), the head mount implantation surgery POD model in this study is not necessarily generalizable to delirium from various causes other than neurosurgery cases.

This study is the first to apply the BSEEG method to POD in animal experiments. We successfully detected the age differences of POD vulnerability in mice quantitatively and chronologically. The data presented here suggests that the BSEEG method may be helpful as a quantitative test for monitoring delirium-like states in mice.

## Acknowledgements

We appreciate the support from Stanford University School of Medicine Veterinary Service Center (VSC).

## Appendixes (Supplementary Materials)

**Supplementary Table 1.**
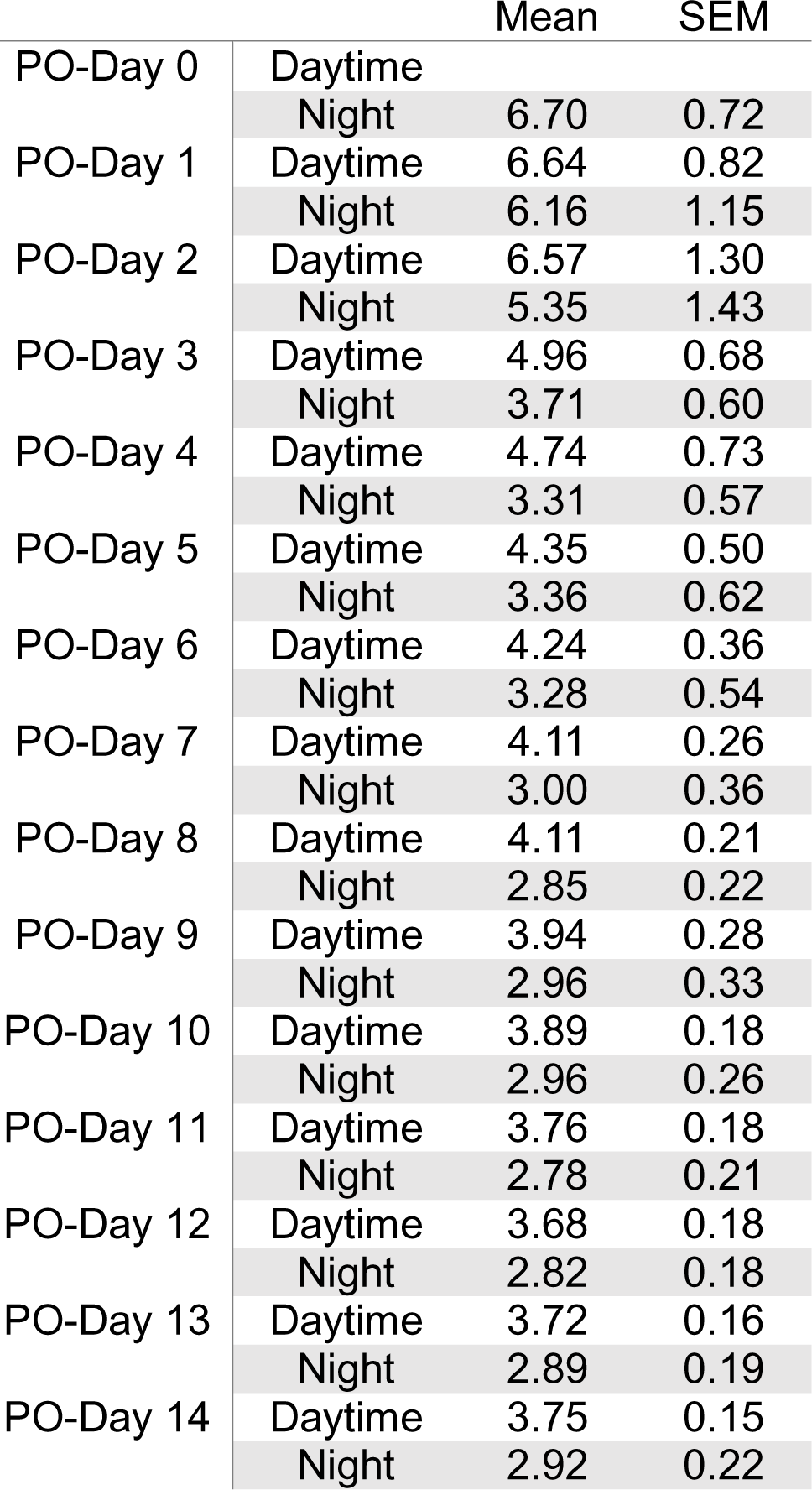
The mean BSEEG score of young mice.

**Supplementary Table 2.** The mean BSEEG score of aged mice.

|  |  | Mean | SEM |
| --- | --- | --- | --- |
| PO-Day 0 | Daytime |  |  |
|  | Night | 6.43 | 0.64 |
| PO-Day 1 | Daytime | 7.54 | 0.68 |
|  | Night | 7.06 | 0.75 |
| PO-Day 2 | Daytime | 7.10 | 0.71 |
|  | Night | 7.21 | 0.93 |
| PO-Day 3 | Daytime | 6.82 | 0.97 |
|  | Night | 6.65 | 1.00 |
| PO-Day 4 | Daytime | 6.12 | 0.96 |
|  | Night | 5.61 | 0.53 |
| PO-Day 5 | Daytime | 6.21 | 0.79 |
|  | Night | 5.84 | 0.75 |
| PO-Day 6 | Daytime | 5.79 | 0.68 |
|  | Night | 5.33 | 0.70 |
| PO-Day 7 | Daytime | 5.58 | 0.66 |
|  | Night | 5.04 | 0.65 |
| PO-Day 8 | Daytime | 5.37 | 0.67 |
|  | Night | 4.32 | 0.37 |
| PO-Day 9 | Daytime | 4.99 | 0.57 |
|  | Night | 4.49 | 0.45 |
| PO-Day 10 | Daytime | 4.81 | 0.56 |
|  | Night | 4.35 | 0.45 |
| PO-Day 11 | Daytime | 4.78 | 0.54 |
|  | Night | 4.39 | 0.46 |
| PO-Day 12 | Daytime | 4.57 | 0.52 |
|  | Night | 4.65 | 0.55 |
| PO-Day 13 | Daytime | 4.66 | 0.52 |
|  | Night | 4.62 | 0.54 |
| PO-Day 14 | Daytime | 4.50 | 0.47 |
|  | Night | 4.65 | 0.51 |

**Supplementary Table 3.**
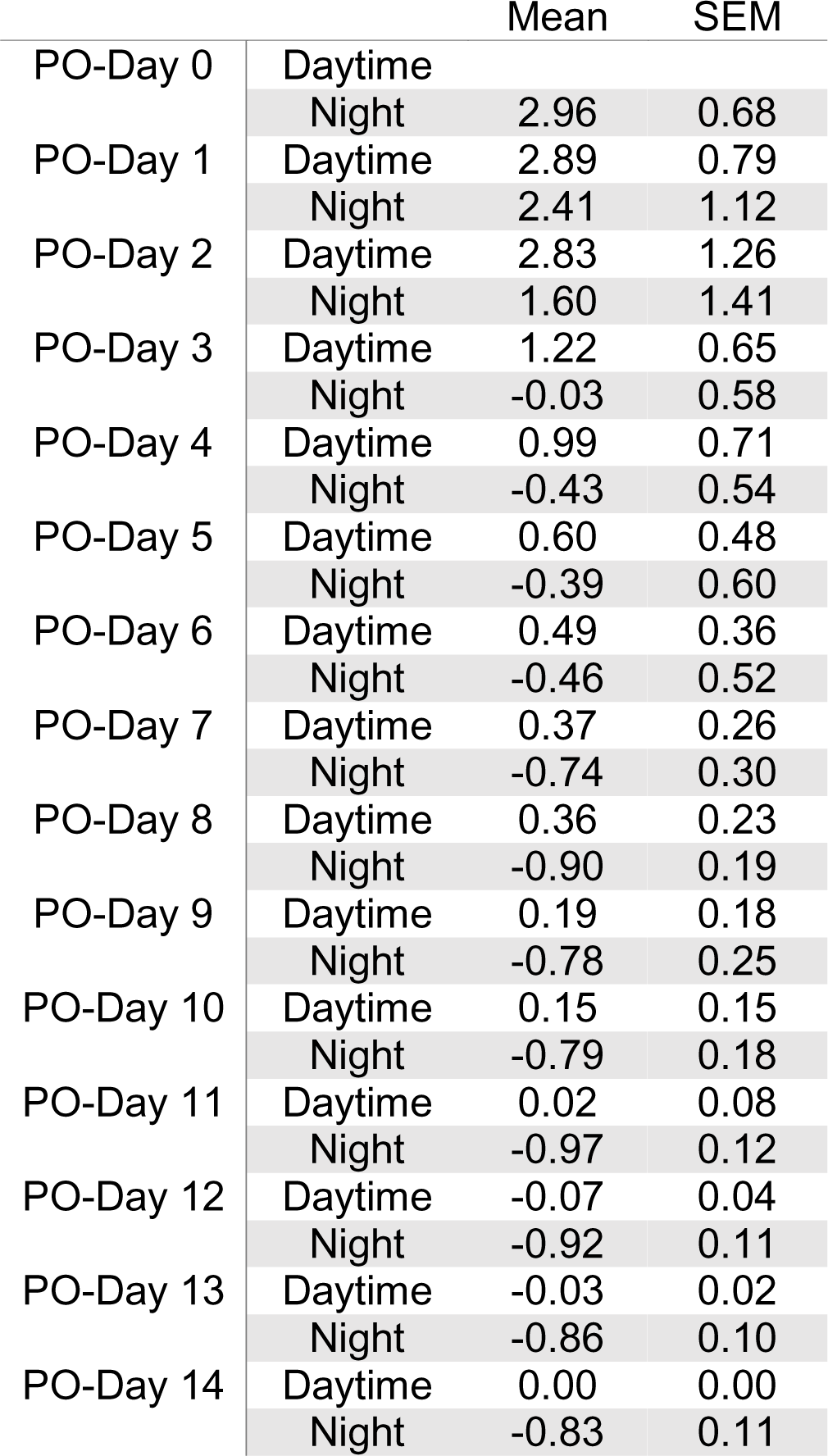
The mean sBSEEG score of young mice.

|  |  | Mean | SEM |
| --- | --- | --- | --- |
| PO-Day 0 | Daytime |  |  |
|  | Night | 2.96 | 0.68 |
| PO-Day 1 | Daytime | 2.89 | 0.79 |
|  | Night | 2.41 | 1.12 |
| PO-Day 2 | Daytime | 2.83 | 1.26 |
|  | Night | 1.60 | 1.41 |
| PO-Day 3 | Daytime | 1.22 | 0.65 |
|  | Night | -0.03 | 0.58 |
| PO-Day 4 | Daytime | 0.99 | 0.71 |
|  | Night | -0.43 | 0.54 |
| PO-Day 5 | Daytime | 0.60 | 0.48 |
|  | Night | -0.39 | 0.60 |
| PO-Day 6 | Daytime | 0.49 | 0.36 |
|  | Night | -0.46 | 0.52 |
| PO-Day 7 | Daytime | 0.37 | 0.26 |
|  | Night | -0.74 | 0.30 |
| PO-Day 8 | Daytime | 0.36 | 0.23 |
|  | Night | -0.90 | 0.19 |
| PO-Day 9 | Daytime | 0.19 | 0.18 |
|  | Night | -0.78 | 0.25 |
| PO-Day 10 | Daytime | 0.15 | 0.15 |
|  | Night | -0.79 | 0.18 |
| PO-Day 11 | Daytime | 0.02 | 0.08 |
|  | Night | -0.97 | 0.12 |
| PO-Day 12 | Daytime | -0.07 | 0.04 |
|  | Night | -0.92 | 0.11 |
| PO-Day 13 | Daytime | -0.03 | 0.02 |
|  | Night | -0.86 | 0.10 |
| PO-Day 14 | Daytime | 0.00 | 0.00 |
|  | Night | -0.83 | 0.11 |

**Supplementary Table 4.**
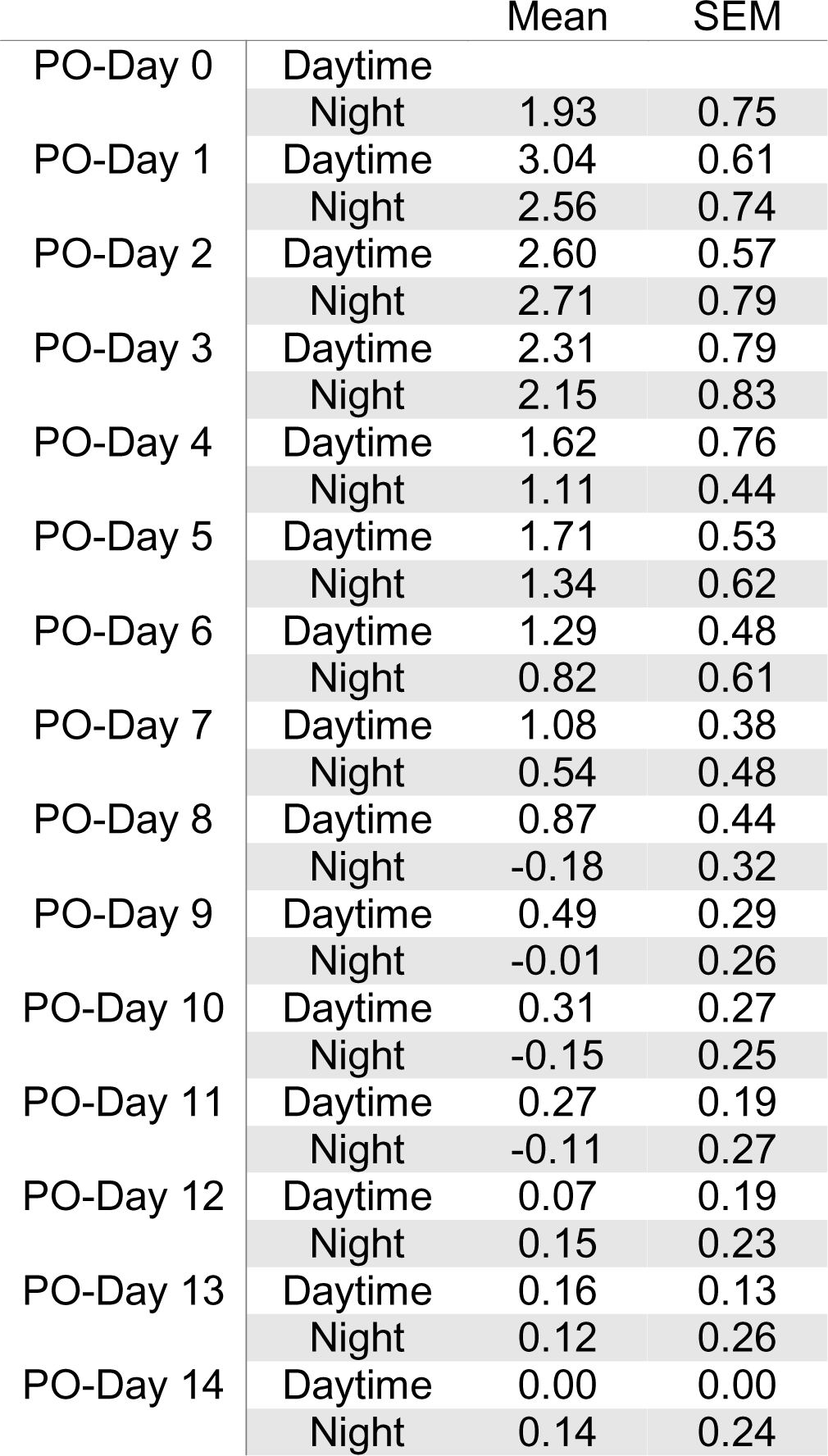
The mean sBSEEG score of aged mice.

**Supplementary Figure 1.**
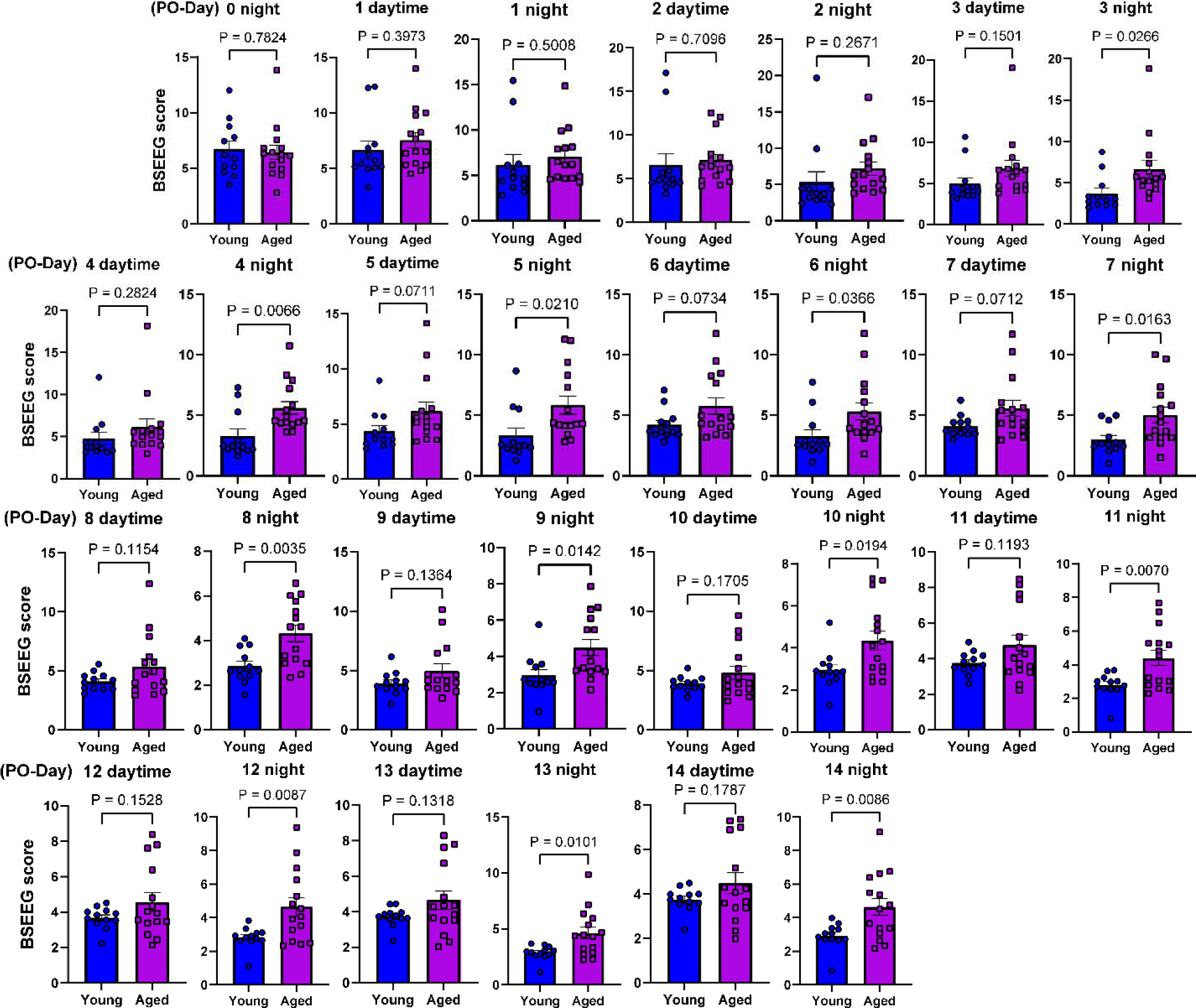
The t-test compares young mice’s BSEEG scores with aged mice’s BSEEG scores.

**Supplementary Figure 2.**
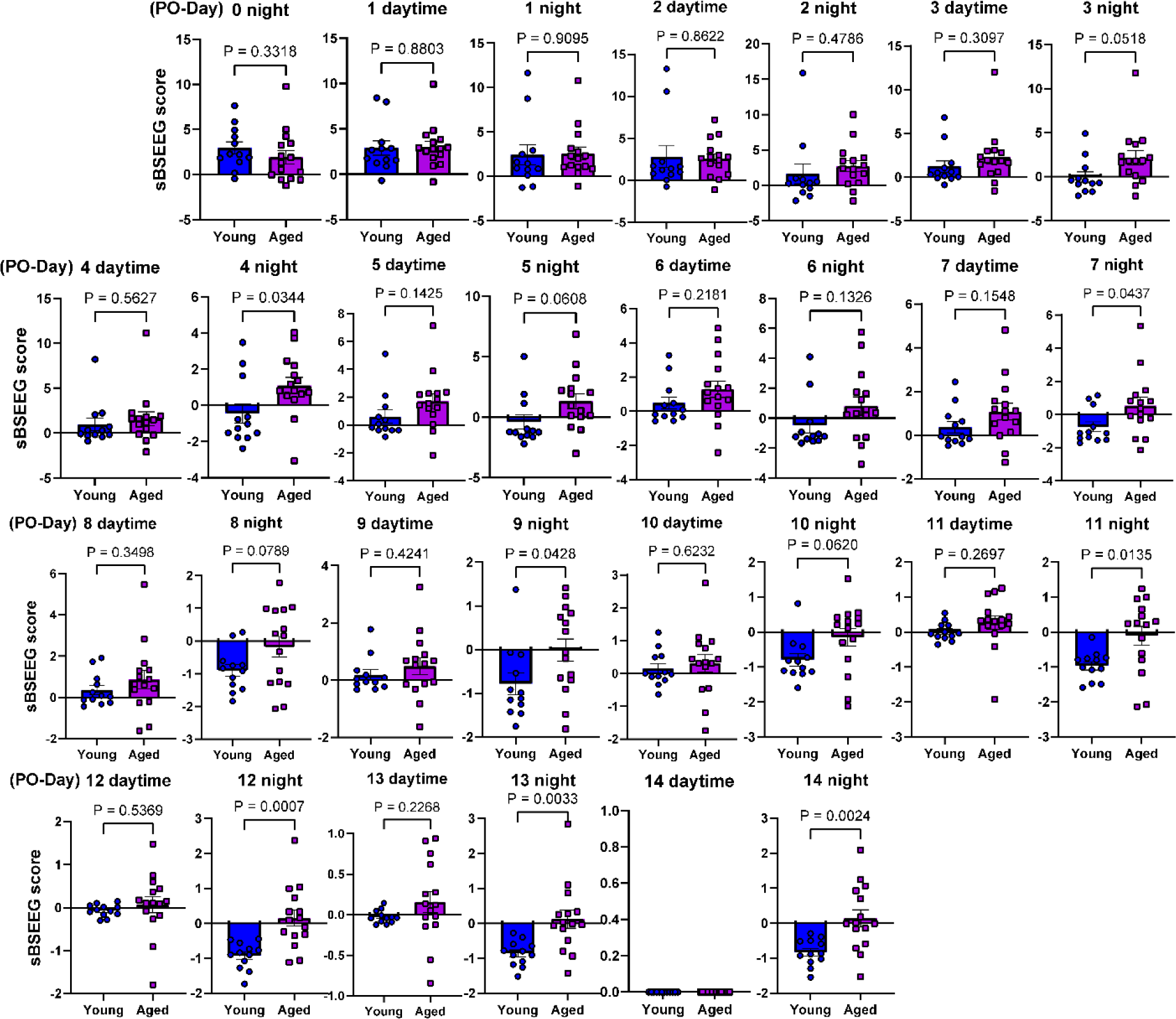
The t-test compares young mice’s sBSEEG scores with aged mice’s sBSEEG scores.

**Supplementary Figure 3.**
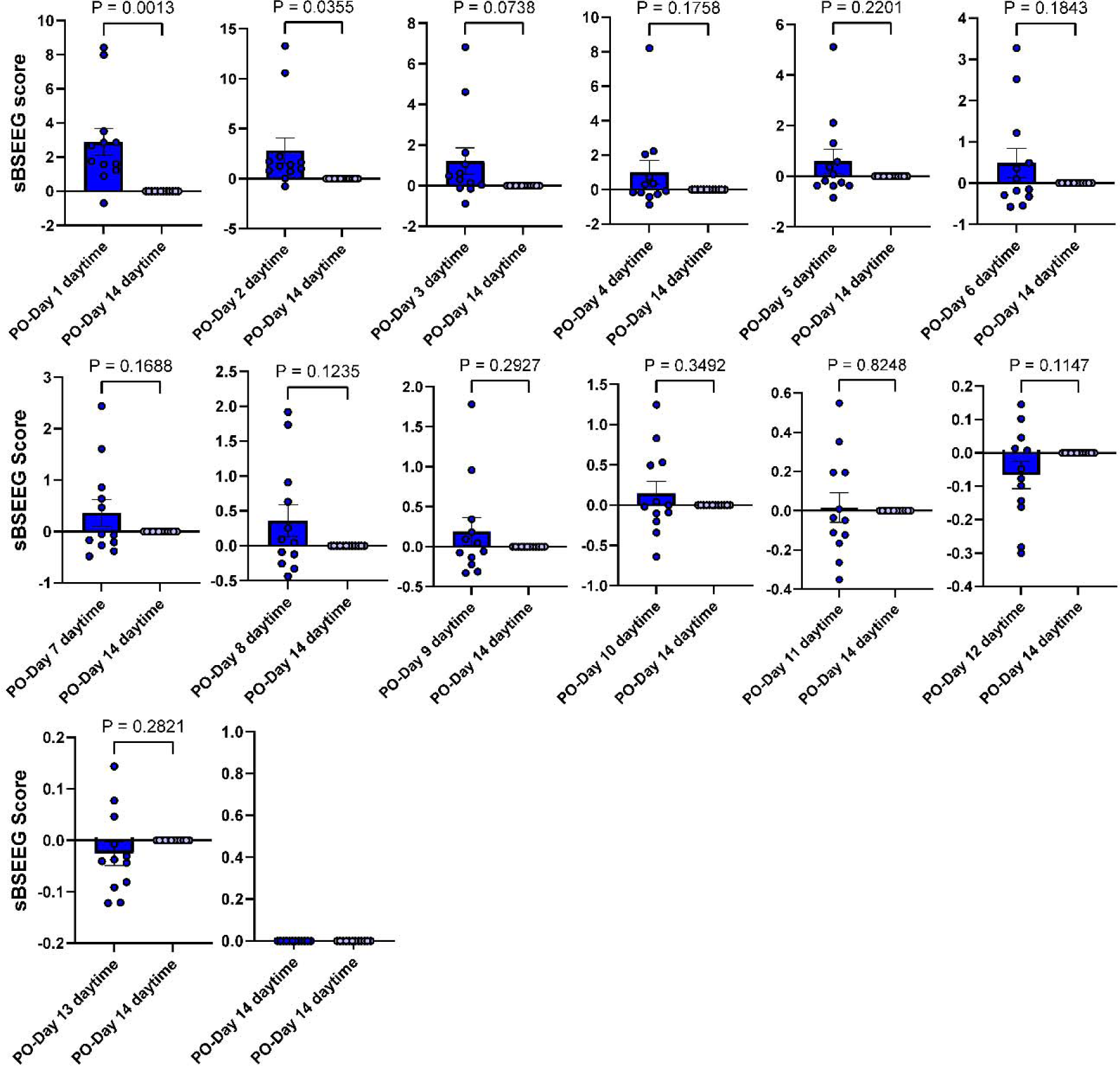
The t-test compares the sBSEEG scores in the daytimes of each PO-Day with the sBSEEG scores as 0 in PO-Day 14 in young mice.

**Supplementary Figure 4.**
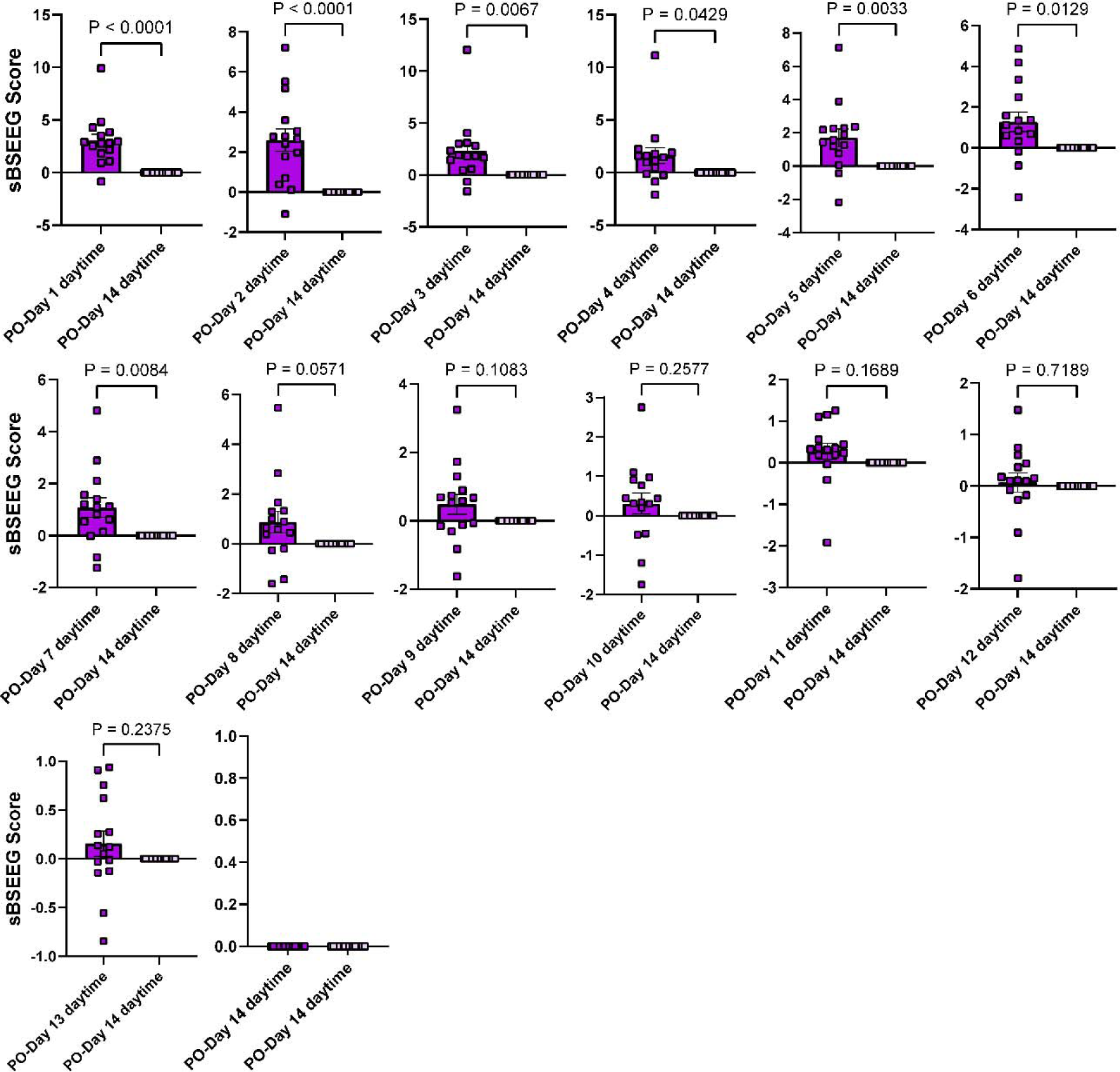
The t-test compares the sBSEEG scores in the daytimes of each PO-Day with the sBSEEG scores as 0 in PO-Day 14 in aged mice.

