## Supplementary figures and images for "The bispectral EEG (BSEEG) method quantifies post-operative delirium-like states in young and aged mice after head mount implantation surgery"

### Supplemental Figure 1

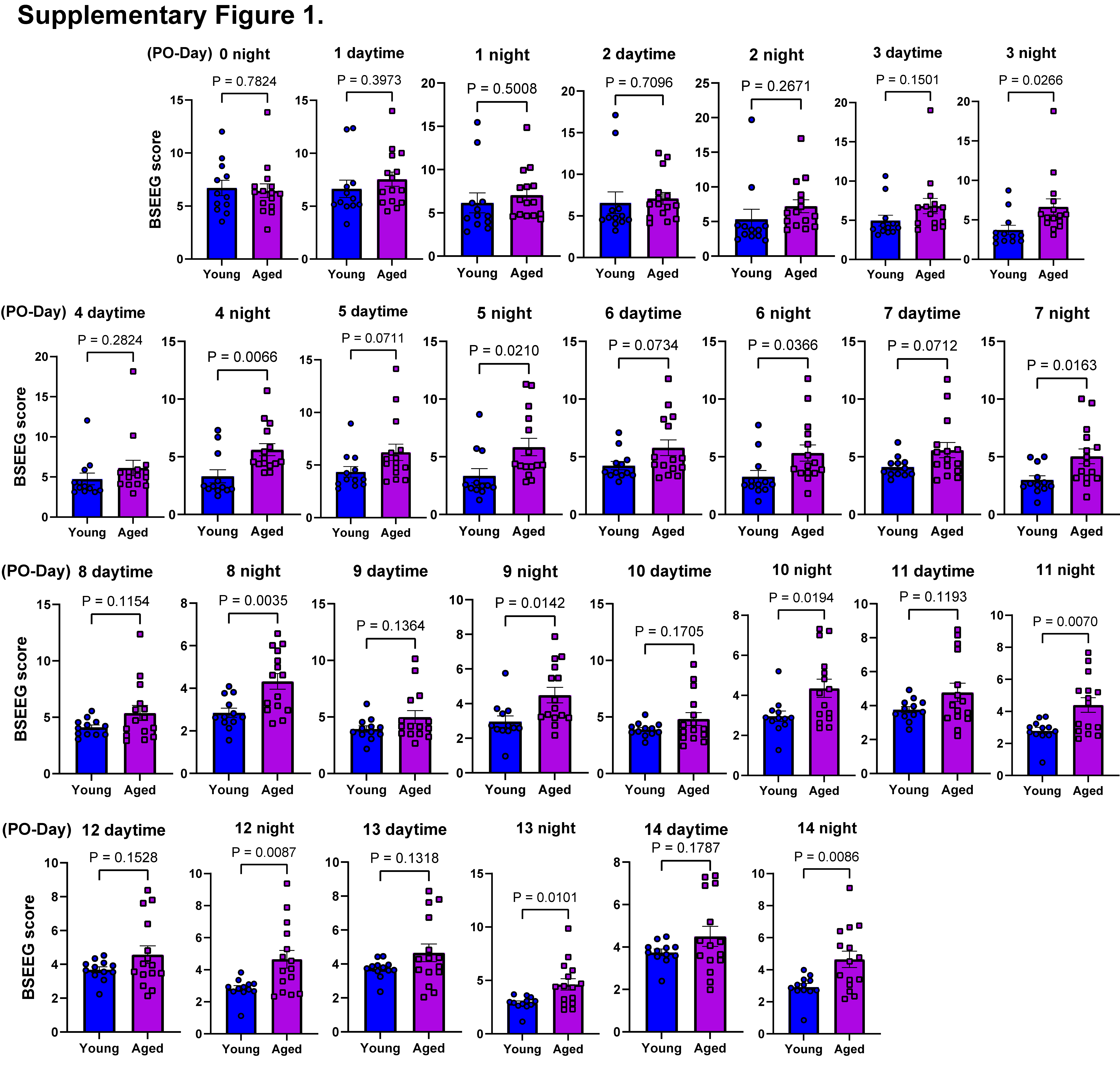

### Supplemental Figure 2

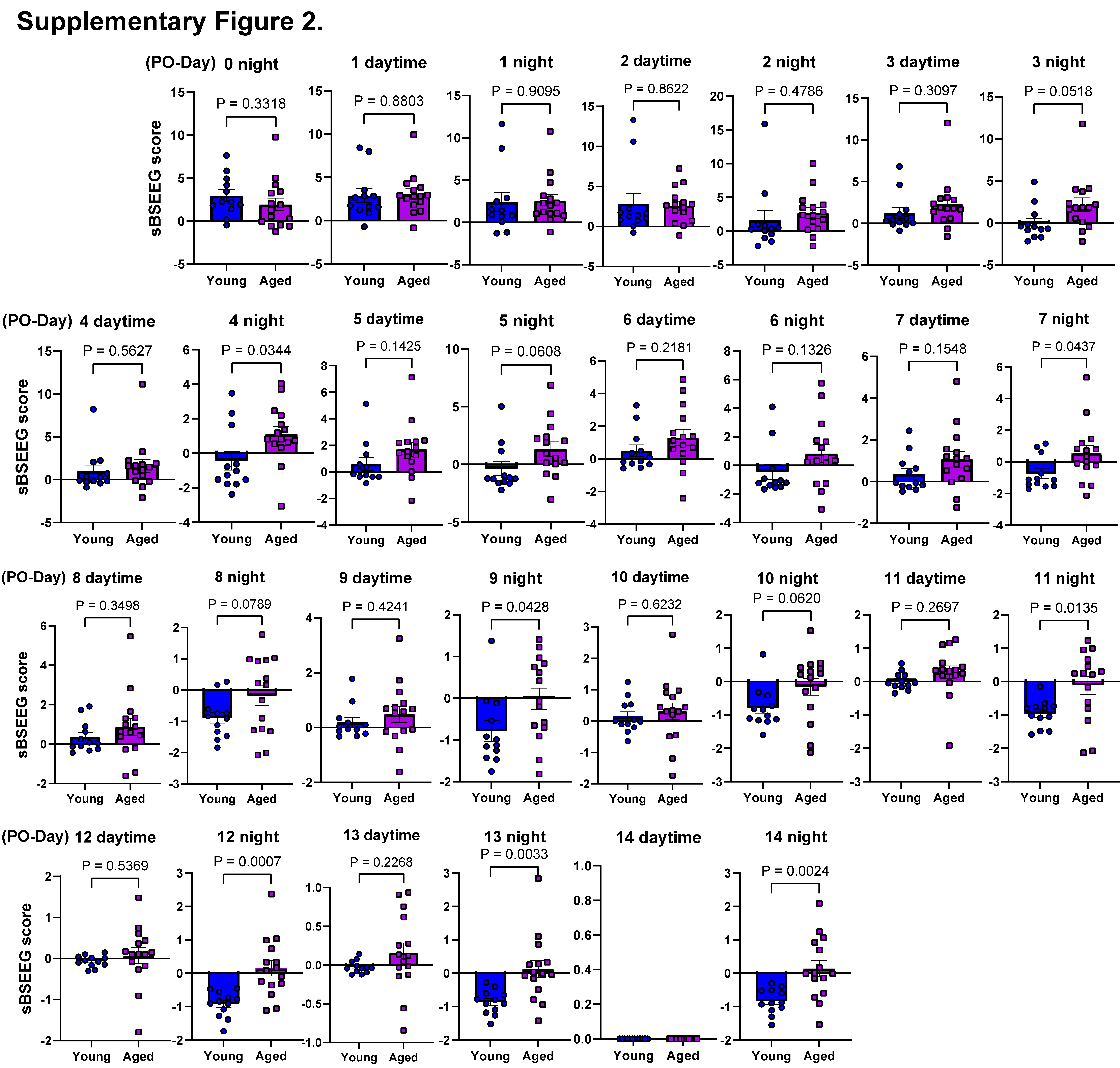

### Supplemental Figure 3

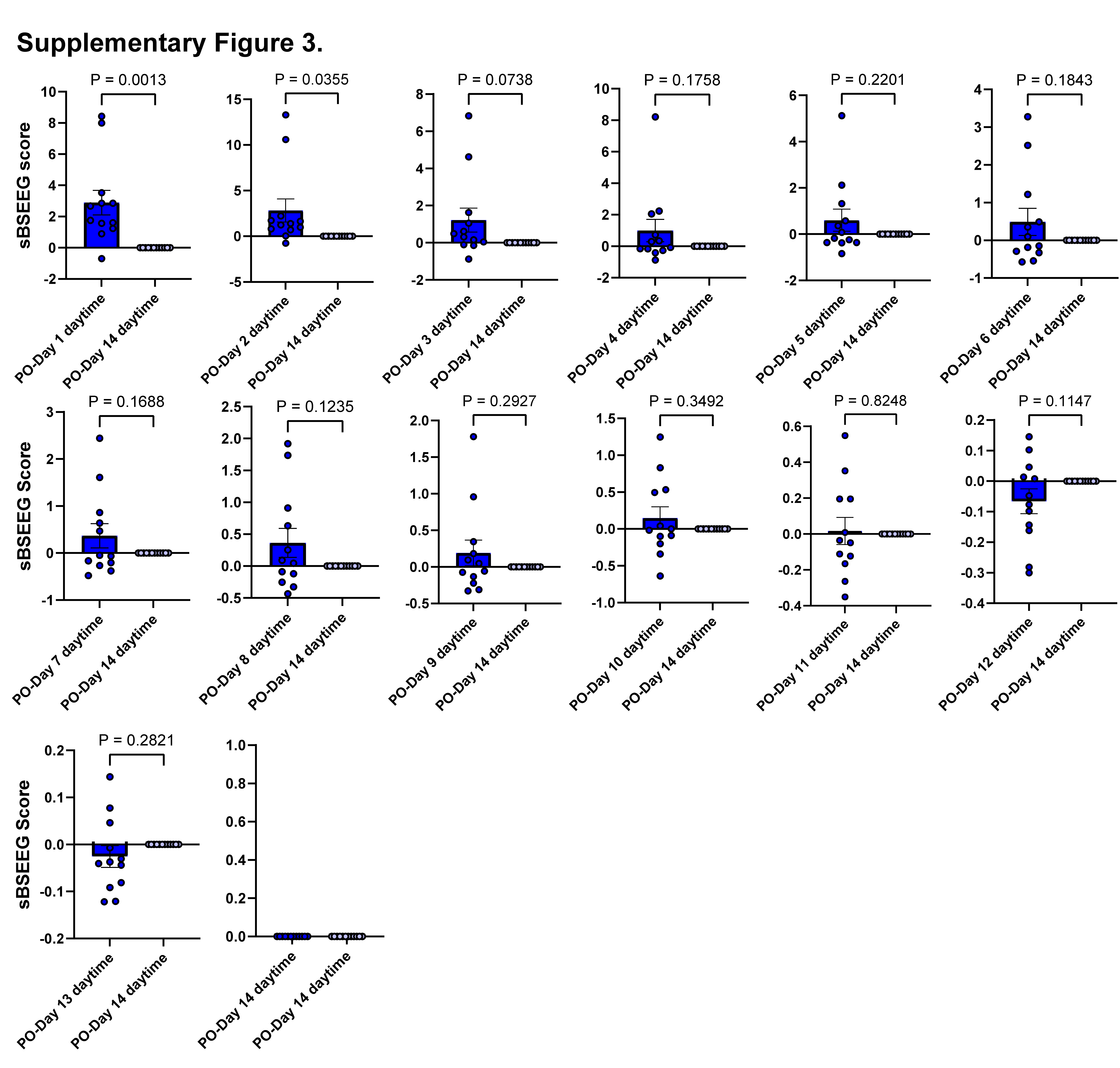

### Supplemental Figure 4

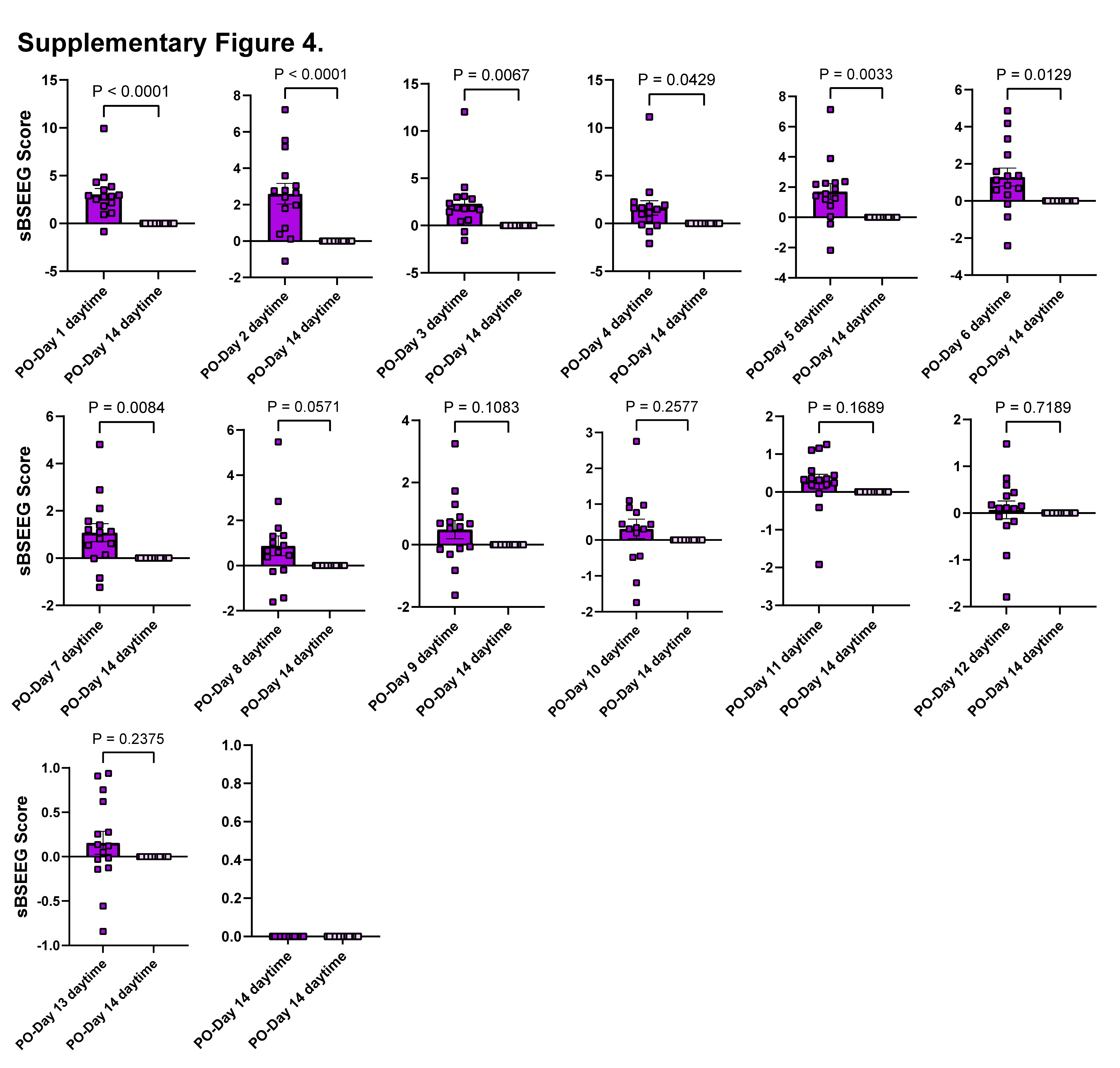
